# The Influence of Parkinson’s Disease and Neurotypical Aging on Cognitive Performance Among Volunteers for an Exercise-based Rehabilitative Intervention

**DOI:** 10.1101/126607

**Authors:** J. Lucas McKay, Ariyana Bozzorg, Joe Nocera, Madeleine E. Hackney

## Abstract

**PURPOSE:** To determine the impact of aging versus combined aging and disease on cognition in older adults with and without Parkinson’s disease (PD) who were volunteers for exercise based rehabilitation research.

**METHODS:** We used a multiple linear regression approach to analyze cognitive outcome measures of rehabilitation volunteers with and without PD.

**RESULTS:** Significant associations were identified between increased age and decreased performance on 8 of the 14 outcomes analyzed after controlling for false discovery rate. Of those 8 outcomes, multivariate regression analyses demonstrated an effect of disease on performance in only 4/8. In all cases, PD was associated with superior, rather than decreased performance after controlling for age. Results were unaffected by sex and education. Post-hoc comparison with available age norms demonstrated that differences between PD and Non-PD volunteers could be primarily attributed to the Non-PD group substantially underperforming versus age norms.

**CONCLUSIONS:** In rehabilitative exercise studies using volunteers, many cognitive domains decline with increasing age, consistent with previous neuropsychological studies without a rehabilitation component. However, older “neurotypical” volunteers may potentially underperform PD volunteers after controlling for age. This may be an important design consideration for rehabilitation studies with cognitive outcomes.

**IMPLICATIONS FOR REHABILITATION:**

- An increasing number of rehabilitation studies incorporate cognitive outcomes.
- Whether the overall cognitive profile of rehabilitation volunteers differs from that of neurotypical aging remains to be established.
- Rehabilitation volunteers with Parkinson’s disease may outperform putatively neurotypical volunteers after controlling for covariates.
- Cognitive impairments associated with PD in neuropsychological studies may not generalize to exercise rehabilitation volunteers.

## INTRODUCTION

In addition to the cardinal motor signs of resting tremor, bradykinesia, rigidity, and postural instability, neuropsychological studies demonstrate that Parkinson’s disease (PD) is also associated with cognitive impairments across multiple domains [1]. At least 25% of PD patients without dementia have comorbid mild cognitive impairment (MCI) [2–4], although standardized diagnostic criteria for PD-MCI were only established relatively recently [5].

Because emerging evidence suggests that exercise-based rehabilitation can benefit cognition in individuals with PD [6–8], cognitive outcome measures are being incorporated in an increasing number of rehabilitation trials. However, it is possible that individuals who are willing to volunteer for extended rehabilitative exercise studies may differ in cognitive profile from individuals who volunteer for crosssectional or observational trials. The cognitive profile of individuals who are willing and able to volunteer for rehabilitation – with or without PD – remains to be established.

Many previous neuropsychological studies have demonstrated that aspects of cognition decline with neurotypical aging [9–13]. This decline may be altered – or may not hold at all – among “neurotypical” volunteers, who may only be inclined to participate due to cognitive or other impairments, e.g., postural instability, that are more prevalent than would be expected of the general population. Similarly, although highly controlled neuropsychological studies have demonstrated broad, mild cognitive impairments in PD [1, 14–16], it is unclear whether this result is reflective of the population of PD patients who are willing and able to volunteer for community-based exercise rehabilitation. Participation in rehabilitation may require PD patients to be comparatively high functioning in order to volunteer [17]. Therefore, it remains unclear whether associations between cognition and neurotypical aging or combined aging and disease will be useful as rehabilitative targets or outcomes.

To open a window into the utility of cognitive outcomes, the purpose of this study was to investigate the relationships between neurotypical aging versus combined aging and neurodegenerative disease on cognition in a sample of older adults with and without PD who had volunteered for exercise-based rehabilitation. We used a multiple linear regression approach to analyze a battery of cognitive outcome measures, including those of visuospatial processing speed, executive function, visuospatial cognition, working memory, incorporation of working memory into motor behavior, mental imagery working memory, attention, and dual tasking. We hypothesized that performance on cognitive outcome measures would be negatively associated both with increasing age and with the presence of PD.

## MATERIALS AND METHODS

### STUDY DESIGN AND DATA SOURCES

All participants provided written informed consent according to protocols approved by the Emory University Institutional Review Board. We performed an observational cross-sectional study using existing baseline data of adults who had volunteered for rehabilitative interventions conducted in 2011-2013. All participants enrolled in Adapted Tango dance rehabilitation [18–25], either without randomization or with the potential for randomization to a health education group. All Non-PD participants were recruited at one of six senior living communities from a large metro area. The senior living communities ranged from high income to very low income and were thus representative of the older adult population residing in senior living communities in a large city in the southern United States. Individuals with PD were recruited from regions in the same metro area, at PD support groups, and educational meetings.

Participants met the following inclusion criteria: no diagnosed neurological conditions other than PD, ability to walk ≥3 meters with or without assistance. Participants with PD met the following additional inclusion criteria: diagnosis of idiopathic “definite PD” [26], and demonstrated response to antiparkinsonian medications.

Beginning with n=116 records, exclusions (n=5 Non-PD, n=3 PD) were applied as follows: < 9 years of formal education / education unknown (n=5), suspected dementia based on Montreal Cognitive Assessment (MoCA) score <16 (n=1) [27, 28], PD onset age < 40 (n=2). Data from participants in one cohort (n=10) to whom the MoCA was not administered were included because of documented typical cognition by study personnel. After applying exclusions, there were 108 individuals available for study. Data from individual instruments (Trail Making test, n=1; Timed Up & Go-Cognitive test, n=1) with strong outlier values were also excluded. Additional outcome measures have been reported previously for a subset of these participants (n=88) [25].

### STUDY VARIABLES

#### Demographic and clinical features of the study population

Demographic and clinical variables included age, sex, years of education, Body Mass Index (BMI), number of comorbidities, and number of prescription medications. Ability to perform Activities of Daily Living (ADLs) was assessed with the Composite Physical Function Index [CPF; 29] and self-reported frequency of weekly trips outside of the home. Global cognitive status was assessed with the MoCA [27,28,30]. Depressive symptoms were assessed with the Beck Depression Inventory-II [BDI-II; 31]. Quality of Life and Fear of Falls were assessed with seven-point Likert scales. Participants with PD were additionally assessed for symptom severity with the motor portion of the Unified Parkinson’s Disease Rating Scale (UPDRS-III) [32]. PD participants were assessed while “ON” antiparkinsonian medications, i.e., a patient-determined optimal time.

#### Primary independent variables and covariates

The primary independent variables were age (years) and dichotomized PD status. Sex and years of education were considered as important covariates because of their strong association with MCI prevalence [4]. Reference coding was used for dichotomous variables to enable interpretation of direction of associations and of the intercept in multivariate analyses. PD status was coded as “1” for PD and “0” for Non-PD, with Non-PD treated as the reference group. Sex was coded as “1” for male and “0” for female, with female as the reference group. Age and years of education were referenced to sample median values.

#### Cognitive outcome measures

Because these data were gathered from individuals participating in exercise studies, outcomes of executive function, mental processing and visuospatial function were included. These cognitive outcomes have been shown to improve after exercise and rehabilitation [33]. Cognitive outcomes included: *Trail Making Test: parts A, B, and B-A*, a test of visual attention, processing speed, and task switching [34–37]; *Reverse Corsi Blocks: Span, Trials, and Product Score*, a test of visuo-spatial short term working memory [38]; *Body Position Spatial Task* (*BPST*)*: Span, Trials, and Product Score*, an adaptation of the Corsi Blocks test incorporating whole-body motion [22]; *Brooks Spatial Task: Percentage Correct*, a test that employs mental imagery to visualize and remember spatial locations [39]. The Brooks Spatial Task employs mental imagery to visualize and remember spatial locations. *Serial Threes: Number Correct and Correct Response Rate* (*CRR*), a simple test of concentration and memory [40]; and *Timed Up & Go-Cognitive* (*TUG-C*)*: Completion Time, Number Correct, and Correct Response Rate* (*CRR*), a measurement of cognitive performance while dual-tasking [40].

**Table 1.** Démographie and clinical features of the study population, stratified by PD status.

| Characteristic | Total Sample<br>(n=108) | PD status |  | Statistic<sup>†</sup> | P |
| --- | --- | --- | --- | --- | --- |
|  |  | PD<br>(n=42) | Non-PD<br>(n=66) |  |  |
| Demographic |  |  |  |  |  |
| Age (years) | 77.6±10.8 | 69.0±8.3 | 83.1±8.3 | 8.59 | <0.01 |
| Sex |  |  |  | 10.48 | <0.01 |
| Female (n) | 62 | 16 | 46 |  |  |
| Male (n) | 46 | 26 | 20 |  |  |
| BMI (kg/m<sup>2</sup>)<sup>a</sup> | 25.1±4.1 | 25.4±3.9 | 24.9±4.3 | 0.61 | 0.54 |
| Comorbidities (#)<sup>a</sup> | 3.2±1.8 | 3.3±1.7 | 3.2±1.8 | 0.27 | 0.79 |
| Prescriptions (#)<sup>b</sup> | 5.0±4.2 | 7.0±5.5 | 3.7±2.2 | 3.57 | <0.01 |
| Cognitive |  |  |  |  |  |
| Education (years) | 15.9±2.4 | 16.9±1.8 | 15.3±2.6 | 3.81 | <0.01 |
| MoCA score (/30)<sup>c</sup> | 23.7±3.2 | 25.2±3.1 | 22.9±3.0 | 3.51 | <0.01 |
| BDI-II (/63)<sup>d</sup> | 8.1±5.8 | 12.2±6.8 | 6.2±4.1 | 4.43 | <0.01 |
| Quality of life (/7) | 5.2±1.1 | 5.1±1.0 | 5.3±1.1 | 1.16 | 0.25 |
| Fear of falls (/7) | 3.0±1.6 | 3.1±1.5 | 3.0±1.6 | 0.43 | 0.67 |
| Activities of daily living |  |  |  |  |  |
| CPF (/24) | 18.8±4.8 | 20.2±4.6 | 17.8±4.7 | 2.55 | 0.01 |
| Weekly trips<sup>a</sup> |  |  |  | 23.6 | <0.01 |
| <1 (n) | 1 | 0 | 1 |  |  |
| 1-2 (n) | 17 | 3 | 14 |  |  |
| 3-4 (n) | 41 | 8 | 33 |  |  |
| daily (n) | 48 | 31 | 17 |  |  |
| Clinical |  |  |  |  |  |
| PD duration (years) |  | 7.0±4.9 |  |  |  |
| UPDRS-III (/108)<sup>e</sup> |  | 28.8±7.0 |  |  |  |
| Hoehn & Yahr (/5)<sup>e</sup> |  |  |  |  |  |
| 3 |  | 9 |  |  |  |
| 2.5 |  | 9 |  |  |  |
| 2 |  | 15 |  |  |  |
| 1.5 |  | 7 |  |  |  |

Abbreviations: PD, Parkinson's disease; SD, Standard Deviation; CPF, Composite Physical Function Index; BMI, Body Mass Index; MoCA, Montreal Cognitive Assessment; BDI-II, Beck Depression Inventory-II; UPDRS-III, Unified Parkinson's Disease Rating Scale, Part III: Motor Exam. <sup>†</sup>All test statistics calculated from independent samples *t*-tests, with the exception of Sex (1-DOF chi-square) and Weekly trips outside the home (3-DOF chi-square).
<sup>a</sup>n=107; <sup>b</sup>n=98; <sup>c</sup>n=99; <sup>d</sup>n=92; <sup>e</sup>n=40.

### ANALYTIC PLAN

#### Descriptive analyses

Descriptive statistics (mean±SD) were calculated for study variables stratified on PD status. Imbalances between the PD and Non-PD groups were evaluated with univariate tests of central tendency (independent sample *t*-tests, chi-square). In cases where the assumption of equal variances was rejected based on Levene’s Test, Satterthwaite’s formula was applied to estimate variances in each group.

#### Bivariate associations between independent variables and cognitive outcome measures

Crude bivariate associations between the two independent variables and each of the cognitive outcome measures were calculated with Pearson’s *r* and validated against point biserial correlation coefficients calculated with special-purpose software [41]. Calculated *r* values were categorized according to cutoff values suggested by Cohen [42].

#### Multivariate associations between age, PD status, and cognitive outcome measures

Linear regression models were used to estimate associations between independent variables age and PD status and each cognitive outcome measure for which a non-negligible value of *r* was identified. Prior to regression analyses, outcome measures were log transformed when necessary in order to decrease right skew. All outcome measures were subsequently standardized to mean value 0 and standard deviation 1. Each outcome measure was then evaluated with linear regression models with age and PD Status entered stepwise. Associations were expressed as β weights and corresponding 95% confidence intervals.

In Model I, age was entered as the independent variable. To control familywise error rate, p values of overall F tests from Model I were corrected using the False Discovery Rate (FDR) method [43]. Outcome measures with FDR-corrected p values ≤ 0.05 in Model I were considered statistically significant and subsequently analyzed with Model II. Remaining outcome measures were not analyzed further.

In Model II, age was entered with PD status as a covariate. PD status was retained as a covariate if two conditions were met: 1) the p value of the overall F test for Model II remained statistically significant (≤ 0.05) after the inclusion of PD Status, and 2) the p value of the t-test for the beta parameter for PD status β_PD_) was ≤ 0.20 in Model II. If both conditions were met, Model II was selected as the “final model” for that outcome measure, and the effect of the inclusion of PD status on the β parameter for age (Δβ_AGE_ was calculated. If not, Model I was selected as the final model for that outcome measure.

#### Controls for education, sex, and post-hoc comparison to normative data

In order to control for imbalances in education and sex, sensitivity of β_AGE_ and β_PD_ in each final model to the inclusion of years of education and sex was also assessed. Post-hoc, outcome measures for which Model II was selected as the final model were converted to age normative data if these were available in the literature in order to compare the Non-PD and PD groups.

## RESULTS

### PARTICIPANT CHARACTERISTICS

Demographic and clinical characteristics of the study population and performance on cognitive outcome measures are summarized in tables 1 and 2, respectively. The entire sample varied in age from 50-95 (PD, 50-82; Non-PD, 59-95). The PD group was younger (14.1 years), more likely to be male (62% vs. 30%), slightly more educated (1.6 years), had better global cognition (2.3 points, MoCA), had less risk of loss of function (2.4 points, CPF), more depressed (6.0 points, BDI-II), and took more prescription medications (7.0 vs. 3.7) than the Non-PD group. The groups were similar in BMI, numbers of comorbidities, and reported generally good quality of life, and mild fear of falling.

**Table 2.** Cognitive outcome measures of the study population, stratified by PD status (n=108).

| Outcome Measure | PD (n=42) |  |  | Non-PD (n=66) |  |  |
| --- | --- | --- | --- | --- | --- | --- |
|  | Mean | SD | Range | Mean | SD | Range |
| Trail Making Test |  |  |  |  |  |  |
| Trail Making A (s)<sup>†</sup> | 43.3<sup>a</sup> | 19.8 | 22.5, 111.8 | 54.6<sup>b</sup> | 22.6 | 26.1, 122.0 |
| Trail Making B (s)<sup>†</sup> | 123.4<sup>a</sup> | 88.6 | 44.4, 300.0 | 151.3<sup>b</sup> | 69.7 | 51.7, 300.0 |
| Trail Making B-A (s)<sup>†</sup> | 80.0<sup>a</sup> | 73.9 | 12.1, 244.1 | 96.7<sup>b</sup> | 59.1 | 13.0, 251.3 |
| Reverse Corsi Blocks |  |  |  |  |  |  |
| Corsi Span | 4.3 | 1.3 | 2, 8 | 3.7<sup>c</sup> | 1.0 | 0, 6 |
| Corsi Trials | 5.8 | 2.2 | 1, 11 | 4.7<sup>c</sup> | 1.5 | 0, 9 |
| Corsi Product Score | 28.2 | 19.9 | 2.0, 88.0 | 18.8<sup>c</sup> | 10.1 | 0.0, 54.0 |
| Body Position Spatial Task |  |  |  |  |  |  |
| BPST Span | 3.7<sup>d</sup> | 0.9 | 2, 5 | 3.3<sup>b</sup> | 0.9 | 1, 5 |
| BPST Trials | 4.3<sup>d</sup> | 1.4 | 2, 8 | 3.5<sup>b</sup> | 1.2 | 2, 7 |
| BPST Product Score | 16.9<sup>d</sup> | 8.8 | 4, 40 | 12.1<sup>b</sup> | 6.9 | 2.0, 35.0 |
| Brooks Spatial Memory Test |  |  |  |  |  |  |
| Brooks Percent Correct | 62.7<sup>e</sup> | 18.1 | 4, 90 | 54.4<sup>f</sup> | 15.8 | 8, 96 |
| Serial Threes |  |  |  |  |  |  |
| Serial Threes #Correct | 7.8<sup>e</sup> | 3.6 | 2.0, 15.3 | 5.5 | 2.6 | 1.7, 14.7 |
| Serial Threes CRR | 0.06<sup>e</sup> | 0.01 | 0.04, 0.07 | 0.06 | 0.01 | 0.04, 0.07 |
| TUG-C |  |  |  |  |  |  |
| TUG-C Completion Time (s)<sup>†</sup> | 14.5<sup>e</sup> | 7.7 | 6.2, 42.6 | 17.7<sup>g</sup> | 7.6 | 7.5, 48.4 |
| TUG-C #Correct | 5.4<sup>e</sup> | 2.8 | 0.0, 14.0 | 4.6<sup>g</sup> | 2.6 | 0.0, 11.0 |
| TUG-C CRR | 0.08<sup>d</sup> | 0.03 | 0.02, 0.18 | 0.06<sup>c</sup> | 0.02 | 0.02, 0.13 |

Abbreviations: PD, Parkinson's disease; SD, Standard Deviation; BPST, Body Position Spatial Task; CRR, Correct Response Rate; TUG-C, Timed Up and Go-Cognitive. <sup>†</sup>Test in which increased score designates poorer performance; otherwise increased scores designate better performance. <sup>a</sup>n=32; <sup>b</sup>n=64; <sup>c</sup>n=63; <sup>d</sup>n=39; <sup>e</sup>n=41; <sup>f</sup>n=60; <sup>g</sup>n=64.

### BIVARIATE ASSOCIATIONS BETWEEN INDEPENDENT VARIABLES AND COGNITIVE OUTCOME MEASURES

Bivariate associations between age, PD status, and cognitive outcomes are summarized in table 3. Negative associations were identified between age and set-switching/executive function, body position spatial cognition, visuospatial processing and span, mental imagery, mental processing/attention and dual cognitive-motor functioning. Associations between age and the majority of outcome measures were of either small (11/15) or medium (2/15) size, with negligible associations identified only for Serial Threes CRR and TUG-C Number Correct.

**Table 3.** Bivariate associations between independent variables and cognitive outcome measures (n=108).

| Outcome Measure | Independent Variable |  |  | PD Status |  |  |
| --- | --- | --- | --- | --- | --- | --- |
|  | Age |  |  |  |  |  |
|  | *r* | *p* | *n* | *r*<sup>†</sup> | *p* | *n* |
| Trail Making Test |  |  |  |  |  |  |
| Trail Making A (s) | 0.15 | 0.14 | 96 | -0.24 | 0.02 | 96 |
| Trail Making B (s) | 0.24 | 0.02 | 96 | -0.17 | 0.10 | 96 |
| Trail Making B-A (s) | 0.23 | 0.02 | 96 | -0.12 | 0.23 | 96 |
| Reverse Corsi Blocks |  |  |  |  |  |  |
| Corsi Span | -0.22 | 0.02 | 105 | 0.27 | <0.01 | 105 |
| Corsi Trials | -0.27 | <0.01 | 105 | 0.29 | <0.01 | 105 |
| Corsi Product Score | -0.27 | <0.01 | 105 | 0.30 | <0.01 | 105 |
| Body Position Spatial Task |  |  |  |  |  |  |
| BPST Span | -0.13 | 0.19 | 103 | 0.22 | 0.03 | 103 |
| BPST Trials | -0.22 | 0.02 | 103 | 0.32 | <0.01 | 103 |
| BPST Product Score | -0.20 | 0.04 | 103 | 0.29 | <0.01 | 103 |
| Brooks Spatial Memory Test |  |  |  |  |  |  |
| Brooks Percent Correct | -0.12 | 0.23 | 101 | 0.24 | 0.02 | 101 |
| Serial Threes |  |  |  |  |  |  |
| Serial Threes #Correct | -0.18 | 0.06 | 107 | 0.40<sup>‡</sup> | <0.01 | 107 |
| Serial Threes CRR | -0.00 | 0.99 | 107 | 0.06 | 0.51 | 107 |
| TUG-C |  |  |  |  |  |  |
| TUG-C Completion Time (s) | 0.38 | <0.01 | 105 | -0.20 | 0.04 | 105 |
| TUG-C #Correct | 0.06 | 0.53 | 105 | 0.14 | 0.14 | 105 |
| TUG-C CRR | -0.48 | <0.01 | 102 | 0.37 | <0.01 | 102 |

Abbreviations: BPST, Body Position Spatial Task; CRR, Correct Response Rate; TUG-C, Timed Up and Go-Cognitive. <sup>†</sup>Pearson's *r* values are identical to point-biserial correlation coefficients unless noted. <sup>‡</sup>Point-biserial correlation coefficient = 0.36.

Associations between PD status and most outcome measures were of either small (10/15) or medium (4/15) size, with negligible associations identified only for Serial Threes CRR. In contrast to negative associations observed between age and outcome measure performance, all non-negligible associations between PD status and outcome measures were positive. Associations between PD status and outcome measures estimated with Pearson’s *r* and the point biserial correlation coefficient were numerically identical to within two significant figures for 14/15 outcome measures and very similar (Serial Threes #Correct; 0.36 vs. 0.40) otherwise. All outcome variables with the exception of Serial Threes CRR were selected for subsequent multivariate analyses.

### MULTIVARIATE ASSOCIATIONS BETWEEN INDEPENDENT VARIABLES AND COGNITIVE OUTCOME MEASURES

After correction for False Discovery Rate, linear regression analyses identified associations between age and decreased performance on 8 of the 14 outcomes analyzed with Model I (table 4). Model I demonstrated significant contributions of age to decreased performance on Trail Making (Part B, *F_1,94_*=11.63, p_FDR_<0.01; B–A (*F_1,94_*=13.51, p_FDR_<0.01), Corsi Blocks (Span, *F_1,103_*=5.45, p_FDR_=0.04; Trials, *F*_1,103_=8.34, p_FDR_<0.01; and Product Score, *F_1,103_*=5.51, p_FDR_=0.04), Body Position Spatial Task (Trials, *F_1,101_*=5.74, p_FDR_=0.04), and Timed Up & Go-Cognitive (Completion Time, *F_1,103_*=26.42, p_FDR_<0.01; CRR, *F_1,100_*=29.24, p_FDR_<0.01).

**Table 4.**
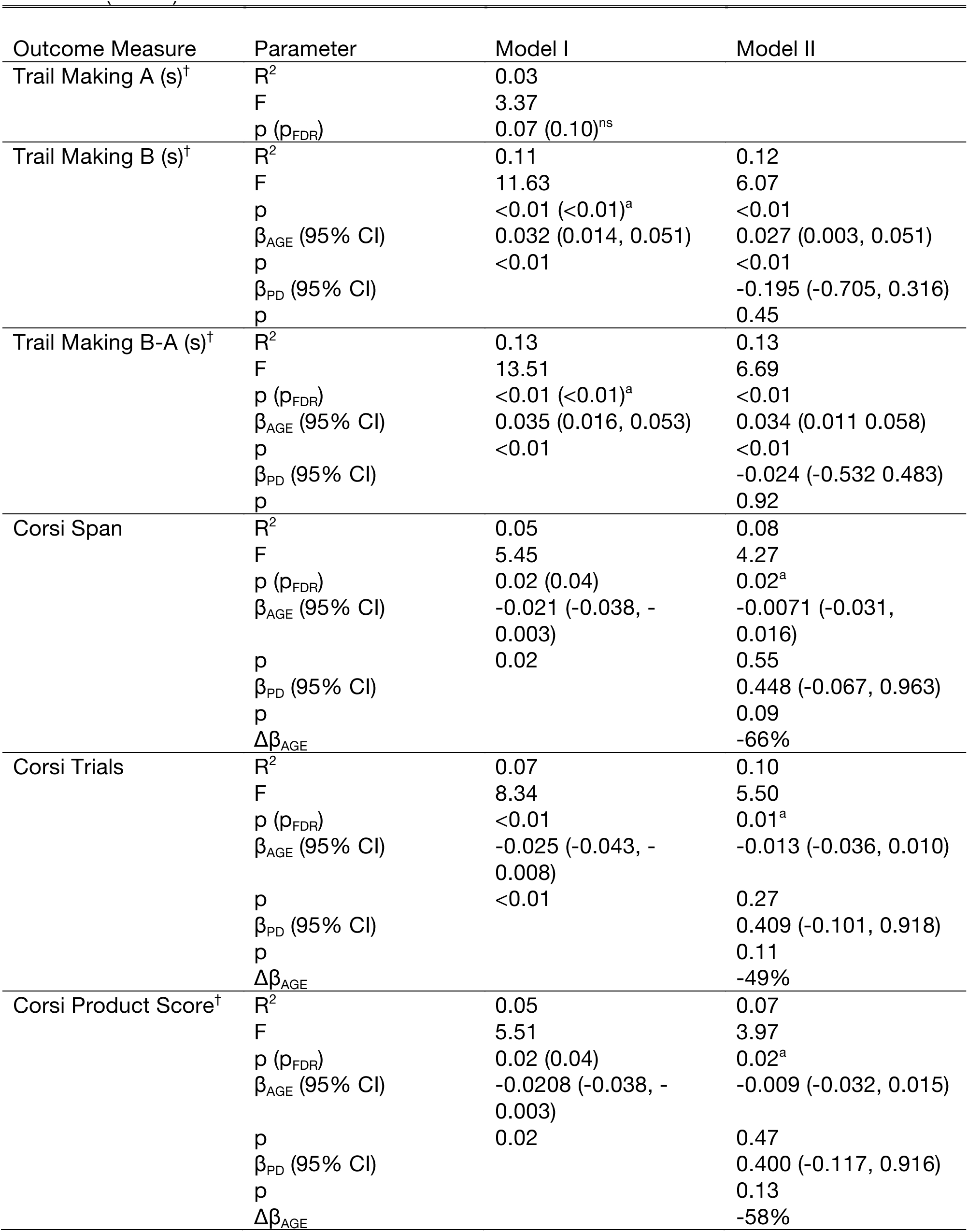
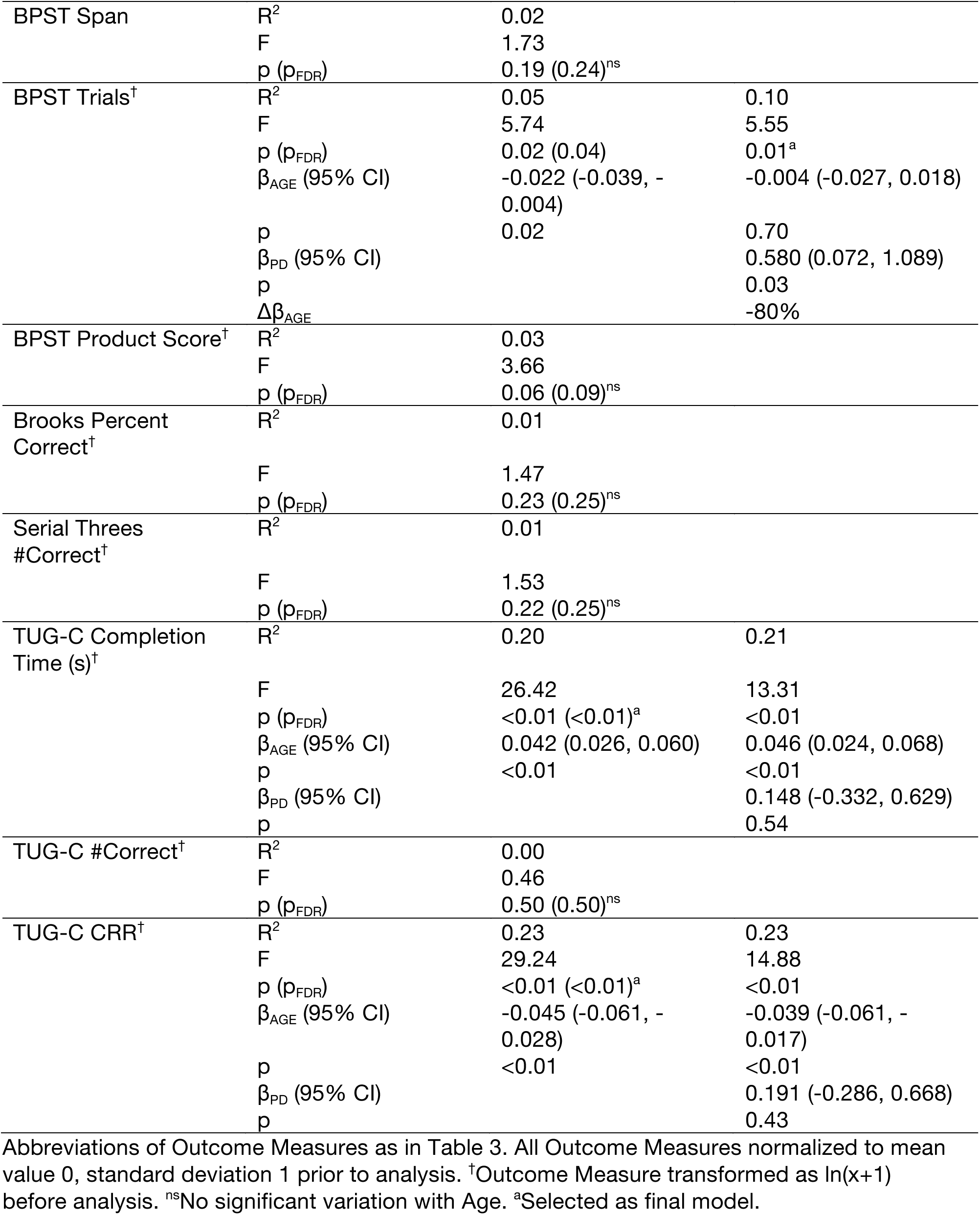
Multivariate associations between independent variables and cognitive outcome measures (n=108).

After accounting for the effects of age in Model I, linear regression analyses with Model II identified associations between PD status and performance on only 4/8 outcome measures examined. All outcome measures for which significant effects of PD status were identified were derived from or adapted from the reverse Corsi blocks paradigm, and in all cases, PD status was associated with increased, rather than decreased, performance after accounting for age (table 4). Including PD status as a covariate resulted in changes to β_AGE_ ≥ 20% for Corsi Blocks (Span, Δβ_AGE_=−66% Model II *F_1,102_*=4.27, p<0.02; Trials, Δβ_AGE_=−49%, *F_1,102_*=5.50, p<0.01; Product Score, Δβ_AGE_=−58%, *F_1,102_*=3.97, p<0.02) and BPST (Trials, Δβ_AGE_=−49%, *F_1,102_*=*5.50*, p<0.01).

### SENSITIVITY OF MULTIVARIATE MODELS TO EDUCATION AND SEX

Neither β_AGE_ nor β_PD_ were affected substantially by the inclusion of years of education and sex as covariates in identified final models. Average changes in regression parameters were 5±5% (range 0-18%) for β_AGE_ and 5±6% (range 0−12%) for β_PD_.

### COMPARISON OF VISUOSPATIAL SHORT-TERM AND WORKING MEMORY PERFORMANCE TO NORMATIVE DATA

In order to interpret the associations between PD status and better performance on cognitive outcome measures identified with Model II, raw scores for Corsi Span, Corsi Trials, and Corsi Product were converted to age-adjusted scores based on normative data of community-dwelling neurotypical older individuals [44]. Mean values and Bonferroni-adjusted (n=6) 95% confidence intervals were calculated for each outcome measure separately for each of the PD and Non-PD groups (figure 1).

**Figure 1.**
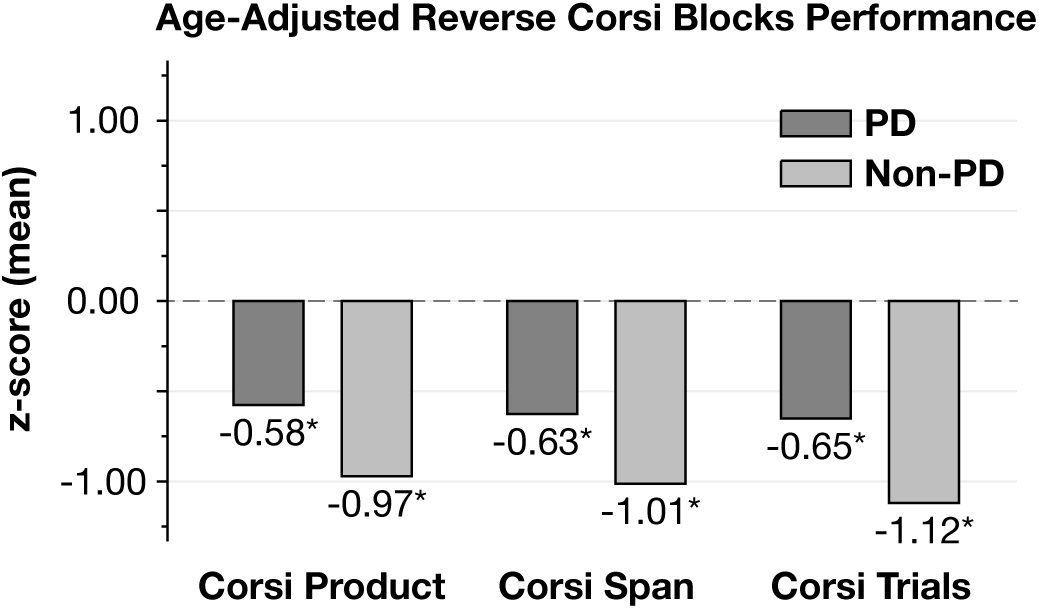
Cognitive outcome measures derived from Reverse Corsi Blocks stratified by PD Status and normalized to scores from neurotypical community-dwelling elderly [44]. *Confidence interval significantly different from zero, p<0.05, after Bonferroni correction for multiple comparisons.

Identified confidence intervals were below zero, the value corresponding to age-normal performance. Confidence intervals calculated for the PD and Non-PD groups were overlapping in all cases. Age-adjusted scores were calculated as follows: Corsi Span, −1.01±1.07, 99.17% CI (−1.38, −0.64) (Non-PD) vs. −0.63±1.16, 99.17% CI (−1.13, −0.13) (PD); Corsi Trials, −1.12±0.91, 99.17% CI (−1.43, −0.81) (Non-PD) vs. −0.65± 1.18, 99.17% CI (−1.56, −0.15) (PD); Corsi Product Score, −0.97±0.75, 99.17% CI (−1.23, − 0.71) (Non-PD) vs. −0.58±1.23, 99.17% CI (−1.10, −0.05) (PD).

No age-normative values for the BPST, an outcome measure developed recently by one of the authors [22], were available in the literature.

## DISCUSSION

Many rehabilitative exercise studies now seek to demonstrate improved cognitive function in PD and other clinical populations as a result of intervention. However, to our knowledge, it is not yet clear whether relationships between aging, disease, and cognition identified in the neuropsychological literature should be assumed to hold among those who volunteer for rehabilitation. In this sample of rehabilitation research volunteers, we found that many cognitive domains declined with increasing age, consistent with previous neuropsychological studies without a rehabilitation component. However, volunteers without PD underperformed volunteers with PD after controlling for age. This may be important to consider in the design of rehabilitation studies with cognitive outcomes.

Similar to the results of previous neuropsychological studies, amongst older rehabilitation volunteers with and without PD, we identified significant associations between increased age and decreased performance on 8 of the 15 cognitive outcomes examined. These included visuospatial processing speed, executive function, visuospatial cognition, working memory, incorporation of working memory into motor behavior, mental imagery working memory, attention, and mental arithmetic performed while walking (dual tasking). Consistent with previously documented declines with age but formerly undocumented in volunteers for exercise research, the strongest associations were observed between age and TUG-C Completion Time and CRR [45], both measures of dual-tasking, and between age and Trail Making B (a test of set-shifting) and Trail Making B-A, which accounts for bradykinesia [35]. These associations were unchanged after controlling for sex and years of education, suggesting that multiple cognitive domains do decline with age among volunteers for rehabilitation.

Surprisingly – and in contrast to the results of neuropsychological studies without a rehabilitation component – after accounting for age, we found negligible differences between the PD and Non-PD groups on four of eight outcome measures that varied significantly with age. When effects of PD were identified, multivariate regression models demonstrated that PD was associated with increased, rather than decreased performance in this sample of rehabilitation volunteers. Bivariate associations revealed that the only exception this trend was in TUG-C completion time, a dual-task test with a substantial whole-body motor component, in which PD was associated with a small impairment, which was not retained as statistically significant in subsequent multivariate models.

These results suggest that the cognitive impairments associated with PD revealed in neuropsychological studies may not generalize to the population of volunteers for exercise rehabilitation designed to enhance motor and cognitive function. It remains unknown whether the mild, broad cognitive impairments identified in neuropsychological studies of PD [1,14,46] will hold in rehabilitation volunteers, because unlike the between-subjects controlled designs often used in neuropsychological studies, rehabilitation studies frequently use repeated-measures designs in a single clinical population, with no data of a well-matched comparison group collected [6–8]. In this sample, post-hoc comparison with available age norms revealed that while both the Non-PD and PD groups performed significantly below age-normal scores, the Non-PD group was substantially more impaired (average z-scores: PD, ≈ −0.60, vs. Non-PD, ≈ −1.00). This is consistent with the interpretation that while the PD group did indeed exhibit some cognitive impairment, the Non-PD group was more impaired – even after accounting for the effects of age.

We speculate that individuals in the PD and Non-PD groups may have experienced different barriers to volunteering, with the result that volunteers in the PD group were less impaired – and that volunteers in the Non-PD group were more impaired – than would be expected for the population in general. As in many community-based rehabilitation studies, exercise classes were held in space donated by senior independent living communities. While all of the individuals in the Non-PD group resided in these centers, the majority (88%) of the individuals in the PD group were required to travel to classes and assessments from their homes. Although in some instances transportation was provided by the study, we speculate that this led to increased barriers to participation among the PD group, and a sample of PD volunteers who were likely higher-functioning than the PD population in general. The potential for study location, and lack of transportation options in underfunded studies, to introduce differential barriers to participation among different study groups may therefore be an important consideration in the design of future rehabilitation trials with cognitive outcomes.

This study has several notable limitations. First, like most geriatric studies, we used a cross-sectional design to gain insight into aging and disease processes. As such, while we did identify effects of age, the actual aging process is certainly more sophisticated than the linear effect modeled here. Second, for parsimony and relevance to exercise rehabilitation, we examined a small subset of cognitive domains to the exclusion of important domains like language, learning and memory, which limits our ability to make inferences about the overall cognitive profile of volunteers for rehabilitation. We were unable to comprehensively normalize the study variables at study onset because normative data remain unavailable or incomplete for many of the cognitive outcome measures used here, many of which have been recently introduced or tailored to PD rehabilitation. For example, although excellent normative data exist for parts A and B of the Trail Making Test [47], we are not aware of any studies that tabulate the difference between parts B and A, which controls for the effects of bradykinesia in parkinsonism [36]. Finally, a worthy comparison could be between individuals with and without PD with similar living arrangements and needs to travel to a study’s interventional program. A design using spouse controls (e.g., [48]) might be appropriate to address this concern.

## CONCLUSION

The results of this cross-sectional study suggest that age-related declines are observed in a broad range of cognitive domains among individuals with and without PD who had volunteered for exercise-based rehabilitation. However, cognitive impairments associated with PD revealed in neuropsychological studies may not generalize to the population of volunteers for rehabilitation.

## ACKNOWLEDGEMENTS

We thank Adrienne Wimberly, Dawa Tsering, Kendra Woodbury, Nathalie Angel, Maria Vasquez and Mylinh Vo for their assistance with assessments and data entry.

